# Start right to end right: authentic open reading frame selection matters

**DOI:** 10.1101/2025.06.10.658836

**Authors:** Mojtaba Bagherian, Georgina Harris, Pratosh Sathishkumar, James P B Lloyd

## Abstract

Accurate annotation of open reading frames (ORFs) is fundamental for understanding gene function and post-transcriptional regulation. A critical but often overlooked aspect of transcriptome annotation is the selection of authentic translation start sites. Many genome annotation pipelines identify the longest possible ORF in alternatively spliced transcripts, using internal methionine codons as putative start sites. However, this computational approach ignores the biological reality that ribosomes select start codons based on sequence context, not ORF length. Here, we demonstrate that this practice leads to systematic misannotation of nonsense-mediated decay (NMD) targets in the Arabidopsis thaliana Araport11 reference transcriptome. Using TranSuite software to identify authentic start codons, we reanalyzed transcriptomic data from an NMD-deficient mutant and found that correct ORF annotation more than doubles the number of identifiable NMD targets with premature termination codons followed by downstream exon junctions, from 203 to 426 transcripts. Furthermore, we show that incorrect ORF annotations can lead to erroneous protein structure predictions, potentially introducing computational artifacts into protein databases. Our findings underscore the importance of biologically informed ORF annotation for accurate assessment of post-transcriptional regulation and proteome prediction, with implications for all eukaryotic genome annotation projects.

## Introduction

Genome sequencing is becoming cheaper and easier, with telomere-to-telomere assemblies becoming routine with third-generation sequencing technologies (Miga *et al*. 2020; Koren *et al*. 2024). However, annotating transcript models of genomes remains a challenge. With RNA-seq, accurate exon-intron boundaries can be determined and whole transcript models can be generated. Multiple transcript models at a genetic locus can be generated via alternative splicing (AS) to increase the transcriptome diversity. But not all alternative transcript isoforms go on to produce protein isoforms. In the case of AS that introduces a premature termination codon (PTC), then the ribosome will terminate early and this will lead to RNA degradation by nonsense-mediated mRNA decay (NMD) (Lloyd 2018), an evolutionarily conserved eukaryotic quality control pathway (Lloyd and Davies 2013; Causier *et al*. 2017). AS-coupled to NMD (AS-NMD) has been reported in animals (Desai *et al*. 2020; Leclair *et al*. 2020; Mironov *et al*. 2023) plants (Stauffer *et al*. 2010; Kalyna *et al*. 2012; Lloyd *et al*. 2018) and fungi (Lareau and Brenner 2015). AS-NMD more than simply NMD removing the products of noisy mis-splicing, but is a highly conserved process (Lareau and Brenner 2015), which is important for the auto- and cross-regulation of many splicing factors(Desai *et al*. 2020; Leclair *et al*. 2020; Mironov *et al*. 2023) and has deep conservation(Lareau and Brenner 2015). Therefore, the regulation of splicing factor level via AS-NMD is an important process to study in health and disease (Leclair *et al*. 2020; Mironov *et al*. 2023).

The commonly reported signals that differentiate a PTC from a normal stop codon are the presence of an exon junction ≥50-55 nucleotides (nt) downstream of the stop codon (50-55 nt rule, or EJC model of NMD), or the presence of a long 3’ UTR (faux 3’ UTR model of NMD) (Lindeboom *et al*. 2016; Lloyd 2018; Lloyd *et al*. 2020; French *et al*. 2020). Studying NMD kinetics at the single-molecule level in human cells has revealed that the exonic sequence downstream of the PTC, PTC to dEJ distance, and number of dEJ all influence NMD efficiency on target transcripts (Hoek *et al*. 2019). However, to accurately predict whether a transcript might be an NMD target, not only do we need accurate structural information for the transcript model (transcriptional start/end sites and exon-intron boundaries), but we also need to know the correct translational start codon and stop codon. An accurate 3’ UTR annotation allows for accurate prediction of NMD-inducing features (Lloyd *et al*. 2018). However, in some genome annotation projects, the algorithms used to find the open reading frame (ORF) within the transcript models simply identifies the longest ORF present in a given transcript. Past publications have highlighted the importance of selecting the biologically authentic ORF rather than simply the longest ORF and the potential consequences (Brown *et al*. 2015; Lloyd *et al*. 2018; Entizne *et al*. 2020), and yet issues still exist (see below) in extant genome annotations. Here we outline why this is a problem for NMD research and protein structural prediction, and then outline a simple solution using freely available software to re-annotate transcriptomes (Entizne *et al*. 2020).

## Results and discussion

For example, the plant model *Arabidopsis thaliana* transcriptome annotation Araport11 (Cheng *et al*. 2017) has numerous examples of the longest ORF being selected for alternatively spliced transcripts (Fig. 1), leading to the appearance of a normal stop codon and an unmodified 3’ UTR (but a large untranslated 5’ UTR). Identifying the longest ORF is computationally simple and logical, but ignores the biological reality: The ribosome cannot predict which ORF is the longest but instead selects the same start codon as if the downstream AS event had not occurred. By annotating the longest ORF through using an internal methionine codon as the new start codon, the 3’ UTR of the AS-coupled NMD target will look normal and the protein will appear as an N-terminal truncation (Fig. 1a and c). In reality, the protein would be a C-terminal truncation and the 3’ UTR would be much longer (Fig. 1b and d) and likely contain at least one downstream exon junction (dEJ). Detection of the altered 3’ UTR can be used to computationally predict a transcript’s NMD-sensitivity (Lindeboom *et al*. 2016; Lloyd 2018; Lloyd *et al*. 2020).

**Fig. 1:**
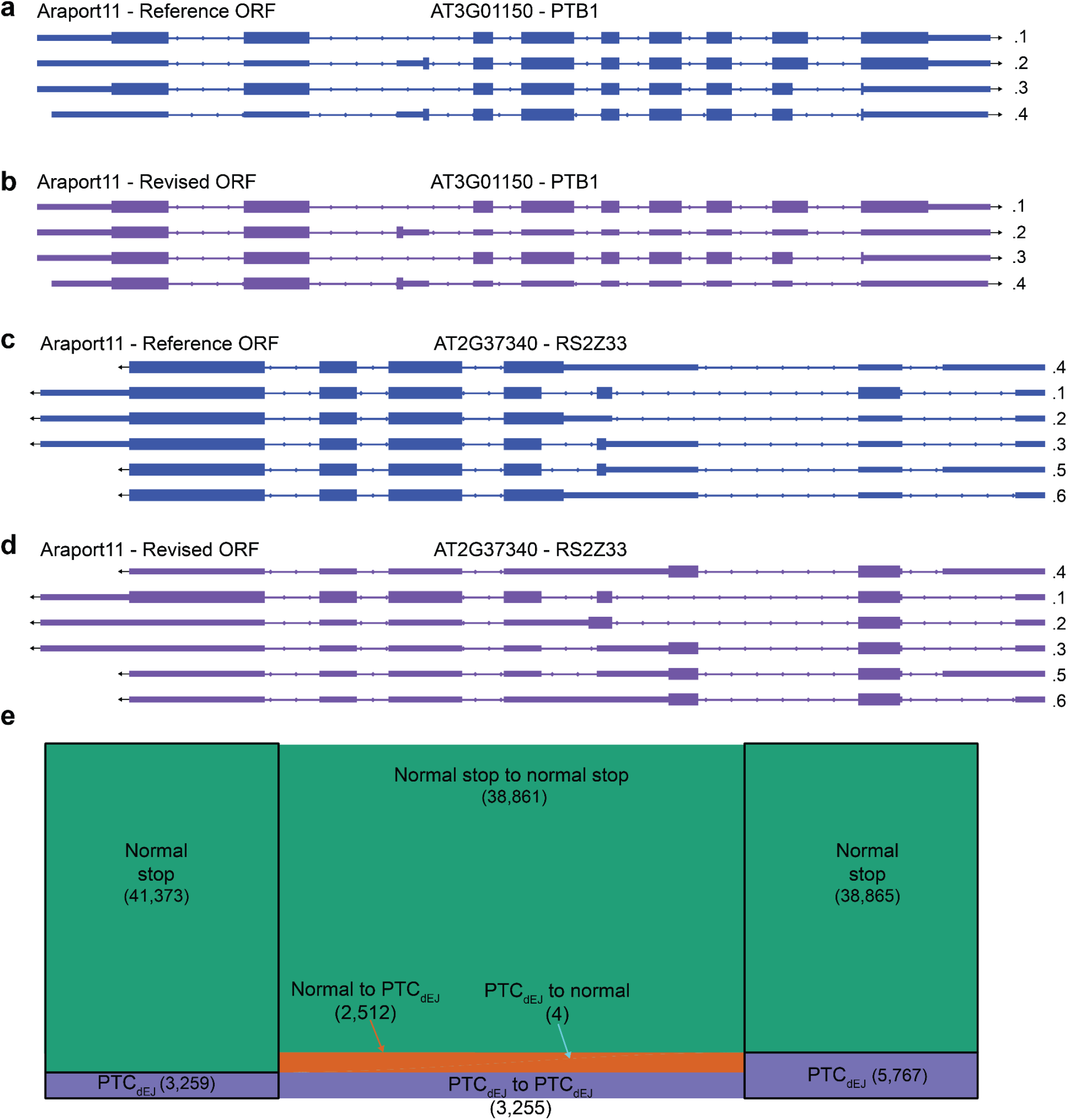
Araport11 transcripts with ORF annotations. **a)** Araport11 - Reference ORF annotations of AT3G01150 (PTB1) gene. **b)** Araport11 - Revised ORF annotations of AT3G01150 (PTB1) gene. **c)** Araport11 - Reference ORF annotations of AT2G37340 (RS2Z33) gene. **d)** Araport11 - Revised ORF annotations of AT2G37340 (RS2Z33) gene. **e)** Change in stop codon status from Araport11 Reference to Araport11 Revised. Normal stop to normal stop (38,861), normal stop to PTC_dEJ_ (2,512), PTC_dEJ_ to normal stop (4), and PTC_dEJ_ to PTC_dEJ_ (3,255).

We took the software TranSuite (Entizne *et al*. 2020), developed to annotate the authentic start codon and stop codon of a transcript, and compared the Araport11 transcriptome’s base annotations (Reference) to the TranSuite’s improved annotations (Revised). The known NMD targets RS2Z33 and PTB1 in Araport11 - Reference ORF annotations have normal 3’ UTRs (Fig. 1a and b), but once annotated with TranSuite, the presence of PTCs as defined by the presence of a dEJ at least 50 nt after the stop codon (PTC_dEJ_) is clear (Fig. 1c and d). Therefore the Revised ORFs generated by TranSuite for these known NMD targets allows for easy detection of the NMD triggering features in these transcripts, while the Reference ORFs appear to have a normal 3’ UTR and would go undetected. Over two thousand transcripts changed from a normal stop codon to a PTC_dEJ_ after TranSuite has been used, while only four go from PTC_dEJ_ to normal stop codon (Fig. 1e). This striking asymmetry likely reflects that TranSuite correctly identifies the authentic start codon that would be used by the ribosome, while the Reference annotation artificially selects downstream methionines that create longer ORFs.

In order to determine the improvement from TranSuite ORF detection on Araport11 transcripts at the transcriptome-wide scale, we compared the Revised ORF with the Reference ORF annotations and the quality of NMD target identification in an NMD-deficient mutant (Drechsel *et al*. 2013). Specifically we looked at transcript isoforms with increased steady state expressed between the wild-type control and the *upf1 upf3* double mutant, which has reduced NMD pathway efficiency (Drechsel *et al*. 2013). After NMD inhibition, direct targets of NMD are expected to increase in expression, however, many transcripts increased and decreased as a result of indirect effect resulting from the loss of NMD. In *A. thaliana*, mutations decreasing NMD activity result in indirect changes in pathogen response (Rayson *et al*. 2012; Riehs-Kearnan *et al*. 2012; Raxwal *et al*. 2020). So we expect only a minority of changing transcripts to be direct targets, which is usually linked to NMD targeting features, such as a long 3’ UTR or downstream exon junction (PTC_dEJ_).

We then compared the fraction of putative NMD targeted transcripts (increased steady state expression) with NMD features with the Araport11 Reference ORF annotations or the Araport11 Revised ORF annotations (as reported by us here using our modified TranSuite (Entizne *et al*. 2020)). We found that only 203 transcripts with increased steady state expression as annotated by Araport11 - Reference ORF annotations had a PTC_dEJ_ (Fig. 2). In contrast, we found that 426 transcripts with increased steady state expression as annotated by Araport11 - Revised ORF annotations had a PTC_dEJ_ (Fig. 2). Not all transcripts with PTC_dEJ_ are expected to be NMD targets, for example, many intron retained transcripts with PTC_dEJ_ are detained in the nucleus (Göhring *et al*. 2014; Boutz *et al*. 2015). However, correct 3’ UTR annotation combined with increased steady state expression in an NMD-deficient mutant is highly indicative of an NMD target. Over two hundred transcripts would have been assumed to be indirect targets of NMD, due to the lack of an abnormal 3’ UTR, if only the Reference ORFs had been considered (Fig. 2a-d). While not all transcripts with a PTC_dEJ_ are expected to be direct targets of NMD, we expect most direct targets to have a PTC_dEJ_, so we predict that the ratio of up/down transcripts with PTC_dEJ_ to be greater than one, and the higher the ratio, the more successful at identifying direct targets of NMD we have been. The ratio of up/down transcripts with PTC_dEJ_ does indeed increase when we move from Araport11 Reference to Revised ORF annotations (Fig. 2c), supporting our notion that improving correct stop codon identification can help in identifying true NMD targets. In contrast, the ratio of up/down transcripts with normal stop codons decreases (Fig. 2d), indicating that many direct targets of NMD were wrongly annotated with normal 3’ UTRs. Together, these results indicate that the TranSuite annotation greatly improves our ability to identify NMD targets. As long-read aided re-annotations of transcript models become more common, such as the *Arabidopsis thaliana* Reference Transcript Dataset 3 (AtRTD3) (Zhang *et al*. 2022), improved ORF annotations are vital to correct interpretation of novel datasets.

**Fig. 2:**
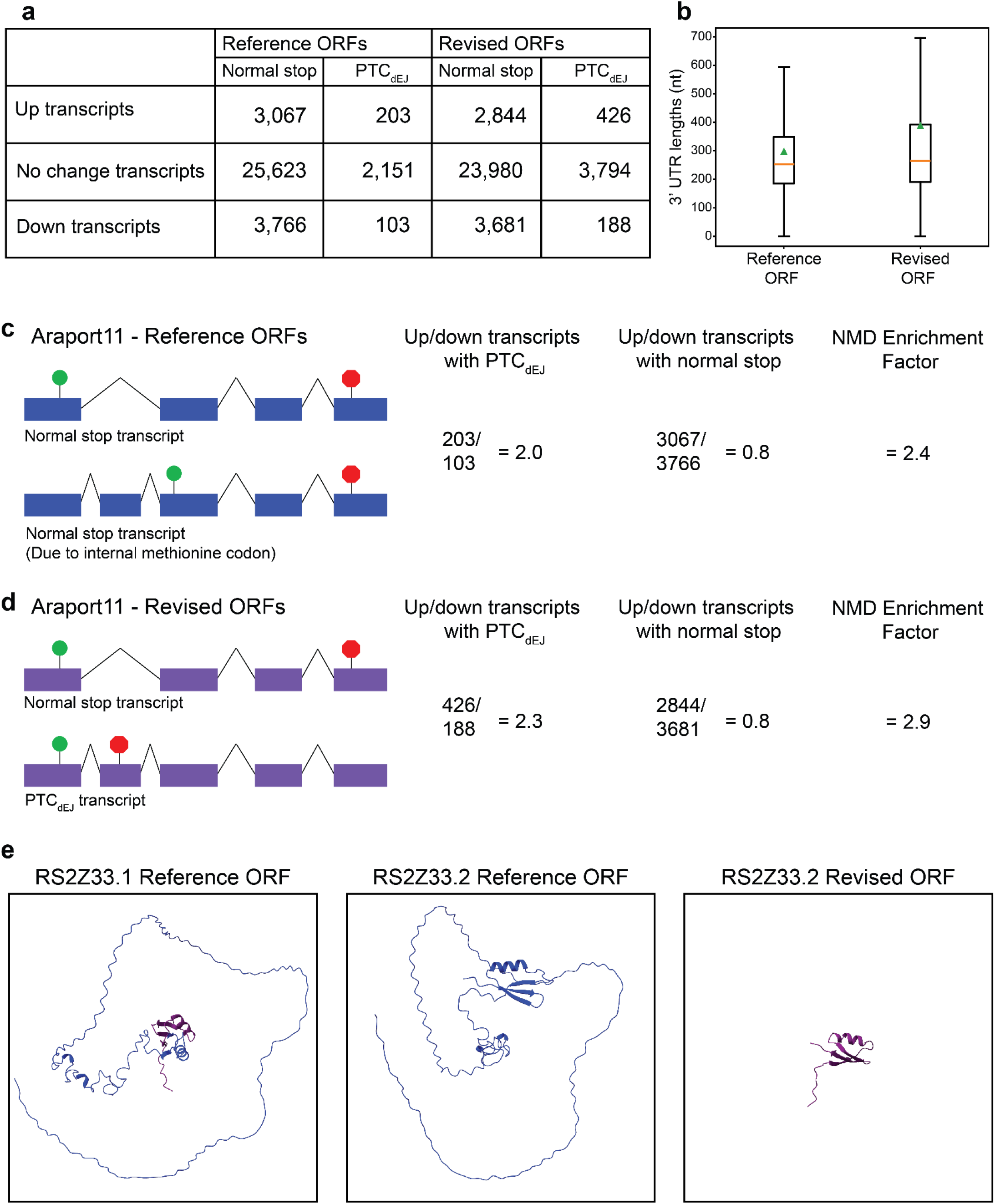
Prediction of authentic start-stop codons improves assessment Araport11 transcripts. **a)** Table indicating the number of up (increased steady state expression), no change, and down (decrease steady state expression) with normal stop codons and PTC_dEJ_ in our transcriptomic re-analysis. **b)** Boxplot (outliers removed) of 3’ UTR lengths (nucleotides; nt) of up (increased steady state expression) transcripts in the NMD-deficient mutant, compared between the Araport11 Reference and Revised ORF annotations. Orange line represents the median. Green triangle represents the mean. Wilcoxon test p = 1.11 × 10^−41^. **c)** Araport11 - Reference ORF annotations and effect on NMD analysis. The number and ratio of up and down transcripts (steady state transcript levels) is shown with either PTC_dEJ_ and normal stop codons. The NMD enrichment factor is the ratio of up/down PTC_dEJ_ transcripts over the ratio of up/down of normal stop transcripts. The higher the number, the higher the enrichment of PTC_dEJ_ changing as predicted based on the EJC model of NMD and not transcripts with normal stop codons. **d)** Araport11 - Revised ORF annotations and effect on NMD analysis. **e)** AlphaFold3-predicted protein structures of RS2Z33 isoforms. Left: RS2Z33.1 (AT2G37340.1) full-length protein. Middle: RS2Z33.2 (AT2G37340.2) Reference ORF - the incorrect long downstream ORF from Araport11. Right: RS2Z33.2 Revised ORF - the correct truncated protein predicted by TranSuite, resulting from intron retention and premature termination of translation.

Computational prediction of protein structure is becoming routine now and automated (Varadi *et al*. 2024), raising the fear that incorrectly annotated ORFs may pollute the databases with the structural predictions of incorrect proteins that are likely to never be translated and have no biological impact on the organism. It has been shown that in the human disease facioscapulohumeral muscular dystrophy, loss of NMD through natural *DUX4* overexpression leads to the production of truncated proteins normally reduced through NMD activity (Campbell *et al*. 2023). As these truncated proteins accumulate, they have a negative effect on cell viability through gain of function activity, as demonstrated by the overexpression of the human SR splicing factor *SRSF3* gene (Campbell *et al*. 2023). If the wrong ORF annotation was used to characterise the predicted truncated protein, misinterpretation would cloud these results. To demonstrate the potential severity of this, we used AlphaFold3 (Abramson *et al*. 2024) to predict the structure of a plant SR splicing factor RS2ZZ33 from this study (Fig. 1b and d). The primary transcript of *RS2ZZ33* (*RS2Z33*.*1*; AT2G37340.1), which encoded the full length protein and is identical between the Araport11 Reference and Revised ORF sets (Fig 2e). When the Araport11 Reference ORF of an intron retained version, *RS2Z33*.*2* (AT2G37340.2), has its structure predicted, we see that it is a long protein with a normal C-terminal but is truncated at the N-terminal, missing this protein fold (Fig 2e). However, when we predict the structure of the Araport11 Revised ORF, we found a small protein with the N-terminal fold only, with the majority of the protein missing due to the early stop codon (Fig 2e). It is unlikely that this truncated protein would be expressed, due to nuclear detention of intron retained transcripts (Göhring *et al*. 2014; Boutz *et al*. 2015), but if translated, this difference in prediction is profound and would have large implications of the estimating the impact of this isoform on molecular outcomes of the cell.

Taken together, we have shown that simply annotating the longest ORF does not reflect the biological reality of transcript anatomy, and that by annotating the authentic start codon, we can see an improvement in fraction of putative NMD identification (Fig. 2). Not only is this important for studying NMD, but it can impact our predictions of protein structures in the proteome. If the reference ORF is used for *in silico* translation and protein structure prediction, the wrong conclusions about the nature of the truncated proteins would be drawn (Fig. 2e). These illegitimately annotated ORFs would lead to computational hallucinations and should be purged from such databases of protein structure. Therefore, not simply annotating the longest ORF of a eukaryotic transcript, but starting with a biologically relevant start codon can have a great benefit for that organism’s genomic resources. For *A. thaliana*, long-read assisted AtRTD3 (Zhang *et al*. 2022) transcriptome already uses TranSuite (Entizne *et al*. 2020) to accurately predict ORFs throughout for accurate prediction of the consequences of AS. When discovering novel splicing events and generating *de novo* transcriptomes (Trapnell *et al*. 2010; Grabherr *et al*. 2011; Pertea *et al*. 2015), we have demonstrated the utility of authentic ORF selection in this process.

## Materials and methods

Genomes: *Arabidopsis thaliana* TAIR10 genome (fasta) with Araport11 transcriptome (GFF3) downloaded from Phytozome. The GFF3 was converted to GTF by the GFFread utility in Cufflinks (Trapnell *et al*. 2010). This was also used to create a transcriptome fasta file from the GTF and genome fasta file.

`gffread -w Athaliana_transcripts.fa -g Athaliana_447_TAIR10.fa Athaliana_447_Araport11.gtf`

The transcriptome fasta file was converted to a Salmon index (kmer setting at 31), Salmon quant was used to to quant transcript abundances (Patro *et al*. 2017). The following settings were used:

`Salmon quant -i A_thaliana_Araport11 -l A -r file.fastq.gz -p 16`

`--validateMappings --fldMean 150 --numBootstraps 100 --seqBias --gcBias -o output`

Previously published RNA-seq data collected from wild-type and *upf1-1 upf3-1* double mutant plants, which have reduced NMD activity (Drechsel *et al*. 2013), were quantified by Salmon (Patro *et al*. 2017), as above. Differential transcript expression was assessed through the R package Sleuth (Pimentel *et al*. 2017). Transcripts were considered differentially expressed with a corrected p-value of < 0.05, and log_2_ fold change > 1 or < 1.

To accurately annotate the authentic start and stop codons of ORFs, rather than just the longest ORF, we used a modified version of TranSuite (Entizne *et al*. 2020). The original version of TranSuite is available here: https://github.com/anonconda/TranSuite. Our modified version of TranSuite is available here: https://github.com/mojtabagherian/TranSuite. Our modified TransFeat module creates a table (CSV) that lists common features related to NMD per transcript, such as distances to the last exon junction downstream of the stop codon and 3’ UTR length. If there was a downstream exon junction and it was at least 50 nucleotides after the stop codon, the transcript isoform is labelled as a Premature Termination Codon downstream Exon-Junction (PTC_dEJ_).

To get correct annotation of Araport11 ORFs as PTC_dEJ_ or not, we filtered out transcript IDs from the GTF that were missing from the TransFix version to allow for a like-for-like comparison and then used our modified TransFeat to identify whether a transcript was PTC_dEJ_ or not. Because many transcripts in Araport11 are annotated with the longest ORF, and internal methionine codons would generate a longer ORF than using the authentic start codon, the stop codon and 3’ UTR does not change in Araport11 - Reference ORF annotations.

Protein sequences were extracted from the default TransFeat module output table (last column, ‘Translation’) generated by TranSuite analysis. The sequences for RS2Z33.1 Reference ORF, RS2Z33.2 Reference ORF, and RS2Z33.2 Revised ORF were submitted to the AlphaFold 3 Server (https://alphafoldserver.com/, Google DeepMind) for structure prediction (Abramson *et al*. 2024). The highest-ranked models (model_0.cif files) were downloaded and visualized using UCSF ChimeraX (v.1.9). Structures were rendered with the RS2Z33.2 Reference ORF shown in blue and the RS2Z33.2 Revised ORF shown in purple to highlight the dramatic structural differences between the longest ORF and the biologically correct ORF predictions.

## Data availability

A Zenodo archive with genomic data is available here: https://doi.org/10.5281/zenodo.15628154. It contains files used to run TranSuite (Athaliana_447_Araport11.gtf, Athaliana_transcripts.fa), intermediary output from TransFind/TransFix (Transfix.gtf), the output from our custom TransFeat (Athaliana_transfeat_splice_junctions.csv). It also contains the modified Araport11 transcriptome (Athaliana_447_Araport11_Reference_ORF.gtf) used as direct input into TransFeat that was used to generate the Reference ORF annotations used in this study (Athaliana_transfeat_splice_junctions_Reference_ORF.csv).

The original version of TranSuite is available here: https://github.com/anonconda/TranSuite. Our modified version of TranSuite is available here: https://github.com/mojtabagherian/TranSuite.

For differential testing, data was downloaded from the NCBI Short Read Archive. SRR584118 (*upf1 upf3* replicate 1), SRR584124 (*upf1 upf3* replicate 2), SRR584115 (wild-type replicate 1), and SRR584121 (wild-type replicate 2).

## Acknowledgment

The computational work of the project was made possible by the High Performance Computing platform *Kaya* at The University of Western Australia maintained by Emily Barker, Chris Bonding and David Gray.

